# Genomic Prediction with Genotype by Environment Interaction Analysis for Kernel Zinc Concentration in Tropical Maize Germplasm

**DOI:** 10.1101/2020.02.24.963090

**Authors:** Edna K. Mageto, Jose Crossa, Paulino Pérez-Rodríguez, Thanda Dhliwayo, Natalia Palacios-Rojas, Michael Lee, Rui Guo, Félix San Vicente, Xuecai Zhang, Vemuri Hindu

## Abstract

Zinc (Zn) deficiency is a major risk factor for human health, affecting about 30% of the world’s population. To study the potential of genomic selection (GS) for maize with increased Zn concentration, an association panel and two doubled haploid (DH) populations were evaluated in three environments. Three genomic prediction models, M (M1: Environment + Line, M2: Environment + Line + Genomic, and M3: Environment + Line + Genomic + Genomic x Environment) incorporating main effects (lines and genomic) and the interaction between genomic and environment (G x E) were assessed to estimate the prediction ability (*r_MP_*) for each model. Two distinct cross-validation (CV) schemes simulating two genomic prediction breeding scenarios were used. CV1 predicts the performance of newly developed lines, whereas CV2 predicts the performance of lines tested in sparse multi-location trials. Predictions for Zn in CV1 ranged from −0.01 to 0.56 for DH1, 0.04 to 0.50 for DH2 and −0.001 to 0.47 for the association panel. For CV2, *r_MP_* values ranged from 0.67 to 0.71 for DH1, 0.40 to 0.56 for DH2 and 0.64 to 0.72 for the association panel. The genomic prediction model which included G x E had the highest average *r_MP_* for both CV1 (0.39 and 0.44) and CV2 (0.71 and 0.51) for the association panel and DH2 population, respectively. These results suggest that GS has potential to accelerate breeding for enhanced kernel Zn concentration by facilitating selection of superior genotypes.

## INTRODUCTION

Malnutrition arising from zinc (Zn) deficiency is a major risk factor for human health affecting nearly 30% of the world’s population (Bouis and Saltzman 2017; Gannon *et al*. 2017). The problem is more prevalent in low-and middle income countries (LMICs), and is highly attributed to lack of access to a balanced diet, reliance on cereal-based diets and ignorance of good nutritional practices (Welch and Graham 2004). Several approaches, such as food fortification, diversification and supplementation have been tried to reduce Zn deficiency. However, in LMICs, these methods have not been entirely successful (Misra *et al*. 2004; Stein 2010).

Breeding maize for increased Zn concentration may offer some relief. The Zn-enriched varieties can be widely accessible, will not require continued investment once developed, and they remain after the initial successful investment and research (Govindan 2011). Recently, maize varieties with 15-36% more Zn were released in Guatemala and Colombia (Listman 2019). Nevertheless, increased breeding efforts are required to develop more Zn-enriched varieties for a diverse range of environments and management practices. Progress toward developing those varieties has mainly relied upon conventional plant breeding approach that is labor-intensive and time-consuming. However, with the recent advances in genomics, new methods for plant breeding such as genomic selection (GS) can be used to identify genotypes with enhanced Zn concentration more efficiently and rapidly.

In a GS breeding scheme, genome-wide DNA markers are used to predict which individuals in a breeding population are most valuable as parents of the next generation (cycle) of offspring (Meuwissen *et al*. 2001; de los Campos *et al*. 2009; Pérez-Rodríguez *et al*. 2012). Kernel Zn concentration is determined at the end of a plant’s life cycle, so GS can enable selection of promising genotypes earlier in the life cycle. This reduces the time and cost of phenotypic evaluation and may increase the genetic gain per unit time and cost (Heslot *et al*. 2015; Manickavelu *et al*. 2017; Arojju *et al*. 2019).

The utility and effectiveness of GS has been examined for many different crop species, marker densities, traits and statistical models and varying levels of prediction accuracy have been achieved (de los Campos *et al*. 2009, 2013; Crossa *et al*. 2010, 2013, 2014; Jarquín *et al*. 2014; Pérez-Rodríguez *et al*. 2015; Zhang *et al*. 2015; Velu *et al*. 2016). Although the number of markers needed for accurate prediction of genotypic values depends on the extent of linkage disequilibrium between markers and QTL (Meuwissen *et al*. 2001), a higher marker density can improve the proportion of genetic variation explained by markers and thus result in higher prediction accuracy (Albrecht *et al*. 2011; Zhao *et al*. 2012; Combs and Bernardo 2013; Liu *et al*.2018). Importantly, higher prediction accuracies have been obtained when genotypes of a population are closely related than when genetically unrelated (Pszczola *et al*. 2012; Combs and Bernardo 2013; Spindel and McCouch 2016).

Initially, GS models and methods were developed for single-environment analyses and they did not consider correlated environmental structures due to genotype by environment (G x E) interactions (Crossa *et al*. 2014). The differential response of genotypes in different environments is a major challenge for breeders and can affect heritability and genotype ranking over environments (Monteverde *et al*. 2018). Multi-environment analysis can model G x E using genetic and residual covariance functions (Burgueño *et al*. 2012), markers and environmental covariates (Jarquín *et al*. 2014), or marker by environment (M x E) interactions (Lopez-Cruz *et al*. 2015). This approach to GS can successfully be used for biofortification breeding of maize because multi-environment testing is routinely used in the development and release of varieties.

Modelling covariance matrices to account for G x E allows the use of information from correlated environments (Burgueño *et al*. 2012). Mixed models that allow the incorporation of a genetic covariance matrix calculated from marker data, rather than assuming independence among genotypes improves the estimation of genetic effects (VanRaden 2008). The benefit of using genetic covariance matrices in G x E mixed models is that the model relates genotypes across locations even when the lines are not present in all locations (Monteverde *et al*. 2018). GS models capable of accounting for multi-environment data have extensively been studied in different crops (Zhang *et al*. 2015; Cuevas *et al*. 2016, 2017; Velu *et al*. 2016; Jarquín *et al*. 2017; Sukumaran *et al*. 2017a; Monteverde *et al*. 2018; Roorkiwal *et al*. 2018). In those studies, incorporating G x E demonstrated a substantial increase in prediction accuracy relative to single-environment analyses.

Kernel Zn has been investigated in several quantitative trait loci (QTL) analyses in maize and each study has reported that Zn concentration is under the control of several loci. The phenotypic variation explained by those loci ranges from 5.9 to 48.8% (Zhou *et al*. 2010; Qin *et al*. 2012; Ŝimić *et al*. 2012; Baxter *et al*. 2013; Jin *et al*. 2013; Zhang *et al*. 2017a; Hindu *et al*. 2018). A Meta-QTL analysis across several of those studies identified regions on chromosome 2 that might be important for kernel Zn concentration (Jin *et al*. 2013). Additionally, genomic regions associated with Zn concentration were recently reported in a genome-wide association study of maize inbreds adapted to the tropics (Hindu *et al*. 2018). Whereas some of the regions were novel, four of the twenty identified were located in already reported QTL intervals. Taken together, the QTLs may be used in a breeding program through marker-assisted selection (MAS) or GS.

A wide array of maize genetic studies has reported considerable effects of G x E interactions for kernel Zn concentration (Oikeh *et al*. 2003, 2004; Long *et al*. 2004; Chakraborti *et al*. 2009; Prasanna *et al*. 2011; Agrawal *et al*. 2012; Guleria *et al*. 2013). However, genotypes with high-Zn concentration have been identified in both tropical and temperate germplasm (Ahmadi *et al*. 1993; Bänziger and Long 2000; Brkic *et al*. 2004; Menkir 2008; Chakraborti *et al*. 2011; Prasanna *et al*. 2011; Hindu *et al*. 2018). Additionally, evaluation procedures for kernel Zn are labor-intensive, expensive and time-consuming (Palacios-Rojas 2018). To the best of our knowledge, no study has examined the predictive ability of GS methods that incorporate G x E for Zn concentration in maize. Within the framework of the reaction norm model (Jarquín *et al*. 2014), the potential of GS for Zn using maize inbreds adapted to tropical environments were assessed. The objectives of this study were; (i) to evaluate the prediction ability for Zn using an association mapping panel and two bi-parental populations evaluated in three tropical environments, (ii) to assess and compare the predictive ability of different GS models, and (iii) to examine the effects of incorporating G x E on prediction accuracy for Zn.

## MATERIALS AND METHODS

### Zinc association mapping (ZAM) panel

The ZAM panel consists of 923 inbreds from maize breeding programs of the International Maize and Wheat Improvement Center (CIMMYT). The panel represents wide genetic diversity for kernel Zn concentration (Hindu *et al*. 2018).

### Bi-parental DH populations

From the ZAM panel, four inbreds with contrasting Zn concentration were selected and used to form two bi-parental (doubled haploid [DH]) populations (Table 1). DH1 was derived from the F1 generation of a mating between CML503, a high-Zn inbred (31.21 μg/g) with CLWN201, a low-Zn inbred (22.62 μg/g). DH2 was derived from the F1 generation of a mating between CML465, another high-Zn inbred (31.55 μg/g) with CML451, a moderate-Zn inbred (27.88 μg/g). DH1 and DH2 were comprised of 112 and 143 inbreds, respectively.

**Table 1.** Pedigree and average concentration of kernel Zn (μg/g) concentration for the parents of the DH populations

| DH population | Pedigree | Parent1 | Parent2 | Zn ($\mu\text{g/g}$) | |
| --- | --- | --- | --- | --- | --- |
|  |  |  |  | Parent1 | Parent2 |
| DH1 | CML503/CLWN201 | CML503 | CLWN201 | 31.21 | 22.62 |
| DH2 | CML 465/CML451 | CML465 | CML451 | 31.55 | 27.88 |

### Experimental design and phenotypic evaluation

### Zinc association mapping (ZAM) panel

The ZAM panel was grown at CIMMYT research stations in Mexico, during the months of June through September and November through March at Agua Fria in 2012 and 2013, and Celaya in 2012. Plot sizes and the experimental designs (Hindu *et al*. 2018).

### Bi-parental DH populations

The DH populations were grown at CIMMYT research stations in Mexico; Celaya in 2014 and Tlaltizapan (18°41’N, 99° 07’ W; 962.5 m asl) in 2015 and 2017. In 2014 and 2015, both populations were evaluated in single-replication trials (Hindu *et al*. 2018). In 2017, a randomized complete block design (RCBD) with two replications was used. The rows were 2.5 m long and 75 cm apart and each genotype was grown in a single row plot. All plots were managed according to the recommended agronomic practices for each environment.

From the ZAM panel and each DH population, four to six plants in each plot were self-pollinated, hand-harvested at physiological maturity, hand-shelled and dried to a moisture content of 12.5%. The bulked kernels from each plot are considered a representative sample and were used in subsequent Zn analyses as described (Hindu et al. 2018).

### Genotypic data

Genomic DNA was extracted from leaf tissues of all inbred lines (ZAM panel and DH populations) using the standard CIMMYT laboratory protocol (CIMMYT, 2005). The samples were genotyped using the genotyping by sequencing (GBS) method at the Institute for Genomic Diversity, Cornell University, USA (Elshire *et al*. 2011; Crossa *et al*. 2013). The restriction enzyme ApeK1 was used to digest DNA, GBS libraries were constructed in 96-plex and sequenced on a single lane of Illumina HISeq2000 flow cell (Elshire *et al*. 2011). To increase the genome coverage and read depth for SNP discovery, raw read data from the sequencing samples were analyzed together with an additional ~30, 000 global maize collections (Zhang *et al*. 2015).

SNP identification was performed using TASSEL 5.0 GBS Discovery Pipeline with B73 (RefGen_v2) as the reference genome (Elshire *et al*. 2011; Glaubitz *et al*. 2014). The source code and the TASSEL GBS discovery pipeline are available at https://www.maizegenetics.net and the SourceForge Tassel project https://sourceforge.net/projects/tassel. For each inbred, the pipeline yielded 955, 690 SNPs which were distributed on the 10 maize chromosomes. After filtering using a minor allele frequency of 0.05 and removing SNPs with more than 10% missing data, 181,889 (ZAM panel) and 170, 798 (bi-parental) SNPs were used for genomic prediction.

### Phenotypic data analysis

For the ZAM panel, broad-sense heritability (*H*^2^) across environments was estimated as:

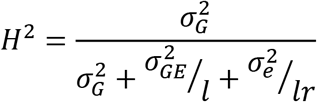

where 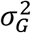 is the variance due to genotype, 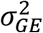 is variance due to genotype x environment, 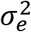 is the error variance, *l* is the number of environments and *r* is the number of replications using multi-environment trial analysis with R (META-R) (Alvarado *et al*. 2016). For the DH populations, variance components based on the genomic relationship matrix were computed using BGLR package as implemented in GBLUP (Pérez and de los Campos 2014). An estimate of narrow-sense heritability (*ĥ*^2^) for each DH population was calculated as:

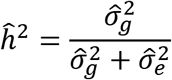

where 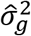 is an estimate of the additive genetic variance and 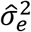 is an estimate of the residual variance.

Correlation coefficients between Zn and environments, descriptive statistics and phenotypic data distribution using boxplots were generated in R (core Team 2018). Line means (genotypic values) for the ZAM panel were estimated as Best Linear Unbiased Estimators (BLUEs) with a random effect for replications nested within each environment. Raw data (values) were used for the DH populations.

### Statistical models

Genomic models used in this study were based on the reaction norm model which models the markers (genomic) by environment interaction (Jarquín *et al*. 2014). This model is an extension of the Genomic Best Linear Unbiased Predictor (GBLUP) random effect model, where the main effects of lines (genotypes), genomic, environments and their interactions are modelled using covariance structures that are functions of marker genotypes and environmental covariates. In this study, environment is the combination of site and year (site-by-year). A brief description of the models is given below.

### M0. Phenotypic baseline model

The phenotypes *y_ij_* are modelled as:

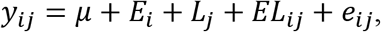

this linear model represents the response of the *j*^th^ (*j* = 1,…, *J*) genotype/line tested in the *i*^th^(*i* = 1,…, *I*) environment 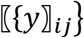 as the sum of an overall mean *μ* plus random environmental main effect 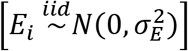, the random genotype main effect 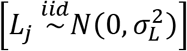, the random interaction between the *j^th^* genotype and the *i*^th^ environment 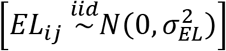 and a random error term 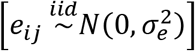. From this linear model, *N*(.,.) denotes a normal random variable, *iid* stands for independent and identically distributed responses and 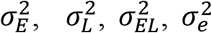 are the variances for environment, genotype, genotype by environment and residual error, respectively. The baseline model does not allow borrowing of information among genotypes because the genotypes were treated as independent outcomes. Thus, models used in this study were derived from the baseline model by subtracting terms or modifying assumptions and/or incorporating genomics/marker information.

### M1. Environment + Line

This model is obtained by retaining the first three components from the baseline model (overall mean, random environment main effect and random line main effect) while their underlying assumptions remain unchanged.

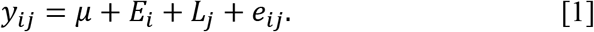

Here environments were considered as site-by-year combinations.

### M2. Environment + Line + Genomic

Another representation of the random main effect of line *L_j_* in the previous model is considering a linear combination between markers and their correspondent marker effects, 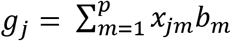, such that

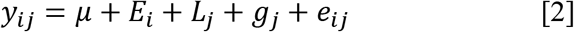

where 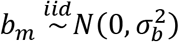 represents the random effect of the *m*^th^ (*m* = 1,…, *p*) marker, *x_jm_* is the genotype of the j^th^ line at the m^th^ marker and 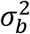 its correspondent variance component. Therefore, **g** = (g_1_,…, g_*J*_)′, is the vector of genetic effects, and follows a normal density with mean zero, and a co-variance matrix 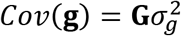 with 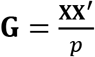 being the genomic relationship matrix (Lopez-Cruz *et al*. 2015) that describes genetic similarities among pairs of individuals. In this model, the line effect *L_j_* is retained to account for imperfect information and model mis-specification because of potential imperfect linkage disequilibrium between markers and quantitative trait loci (QTLs).

### M3. Environment + Line + Genomic + Genomic × Environment

This model accounts for the effects of lines *L_j_*, of markers (genomic) *g_j_*, of environments (*E_i_*) and the interaction between markers (genomic) and the environment (*Eg_ij_*). The model includes the interaction between markers (genomics) and the environment via co-variance structure (Jarquín *et al*. 2014). The model is as follows:

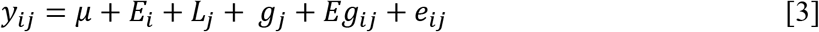

Where *Eg_ij_* is the interaction between the genetic value of the *i^th^* genotype in the *j^th^* environment and 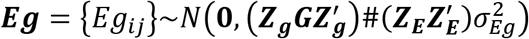, where ***Z***_*g*_ and ***Z***_*E*_ are the correspondent incidence matrices for the effects of genetic values of genotypes and environments, respectively, 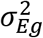 is the variance component of ***Eg*** and # denotes the Hadamard product (element-to-element product) between two matrices.

### Prediction accuracy assessment using cross-validation

Two distinct cross-validation schemes that mimic prediction problems that breeders may face when performing genomic prediction were used (Burgueño *et al*. 2012). One random cross-validation (CV1) evaluates the prediction ability of models when a set of lines have not been evaluated in any environment (prediction of newly developed lines). In CV1, predictions are entirely based on phenotypic records of genetically related lines. The second cross-validation (CV2) is related to incomplete field trials also known as sparse testing, in which some lines are observed in some environments but not in others. In CV2, the goal is to predict the performance of lines in environments where they have not yet been observed. Thus, information from related lines and the correlated environments is used, and prediction assessment can benefit from borrowing information between lines within an environment, between lines across environments and among correlated environments.

In CV1 and CV2, a fivefold cross-validation scheme was used to generate the training and validation sets to assess the prediction ability for Zn within the ZAM panel and each DH population. The data were randomly divided into five subsets, with 80% of the lines assigned to the training set and 20% assigned to the validation set. Four subsets were combined to form the training set, and the remaining subset was used as the validation set. Permutation of five subsets taken one at a time led to five training and validation data sets. The procedure was repeated 20 times and a total of 100 runs were performed in each population. The average value of the correlations between the phenotype and the genomic estimated breeding values (GEBVs) from 100 runs was calculated for the ZAM panel, and each DH population for Zn in each environment and was defined as the prediction ability (*r_MP_*).

### Data availability

All models were fitted in R (core Team 2018) using the BGLR package (Pérez and de los Campos 2014). All phenotypic and genomic data can be downloaded from the link: http://hdl.handle.net/11529/10548331

## RESULTS

### Descriptive statistics

Mean values of kernel Zn concentration were estimated for each environment and across environments (Tables 2A and 2B). For the ZAM panel, kernel Zn ranged from 14.76 to 39.80 μg/g in Celaya 2012, 15.16 to 42.52 μg/g and 17.05 to 46.52 μg/g in Agua Fria 2012 and 2013, respectively (Figure 1). The highest mean (29.53 μg/g) for Zn was observed in Agua Fria 2013. DH1 had Zn values ranging from 16.00 to 48.00 μg/g in Celaya 2012, 16.00 to 35.00 μg/g in Tlaltizapan 2015 and 15.50 to 39.00 μg/g in Tlaltizapan 2017, while the respective values for DH 2 were 17.70 to 43.14 μg/g, 15.60 to 37.80 μg/g and 14.70 to 37.60 μg/g (Figures 2A and 2B). The highest means for Zn were observed in Celaya 2014 (25.38 μg/g) and 2017 (27.96 μg/g) for DH1 and DH2, respectively (Table 2B). Across environments, heritability 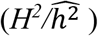 estimates were 0.85, 0.83 and 0.76 for the ZAM panel, DH1 and DH2, respectively (Tables 2A and 2B). There were significant positive correlations between environments for Zn (Table 3), accounting for the moderate to high heritability estimates.

**Figure 1.**
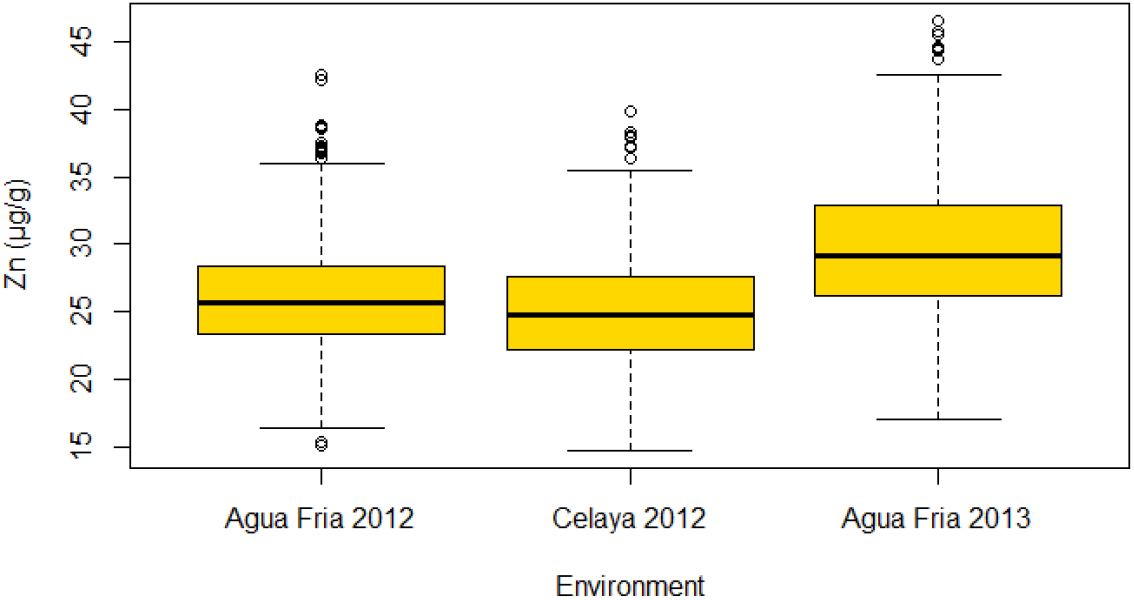
Box plots for kernel Zn (μg/g) in the ZAM panel in three environments (Agua Fria, 2012, Celaya, 2012 and Agua Fria, 2013)

**Figure 2.**
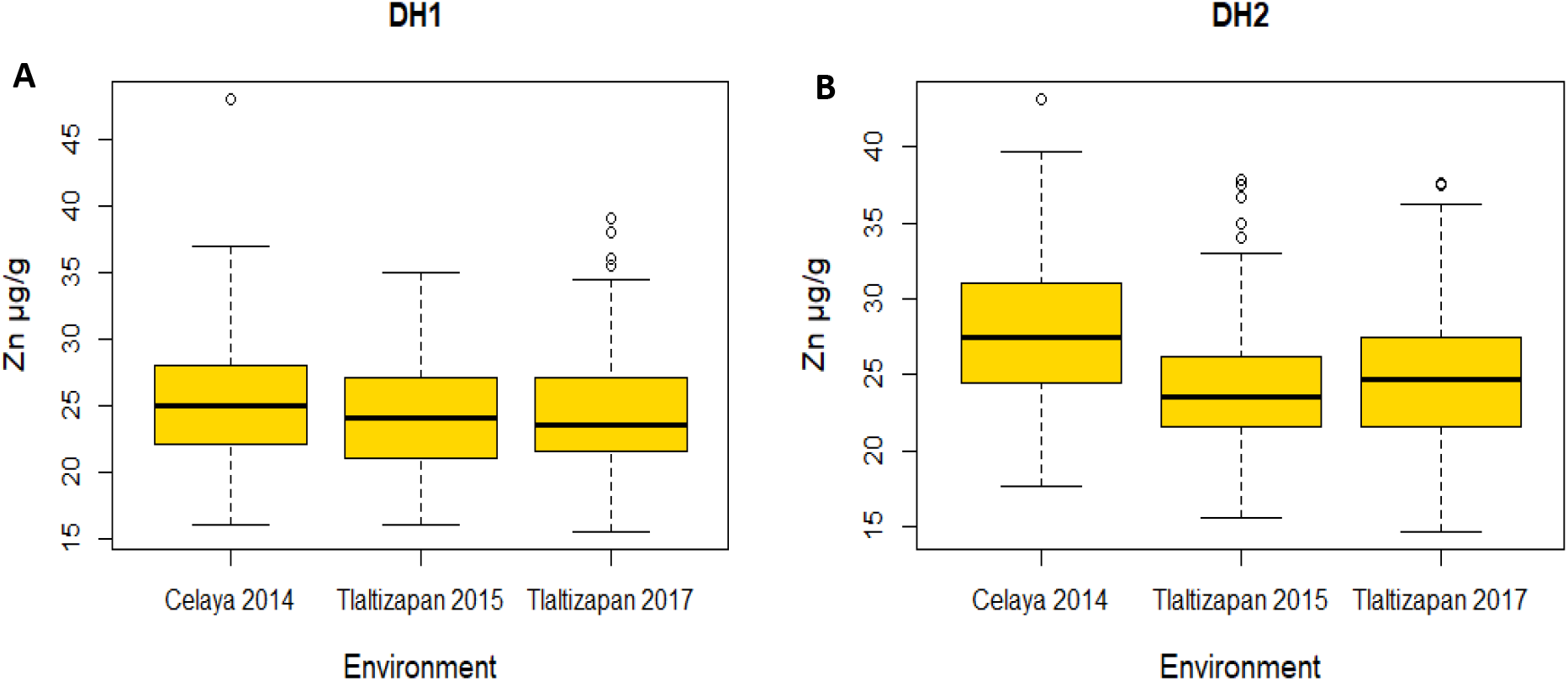
Box plots for kernel Zn (μg/g) for (A) DH1 and (B) DH2 in three environments (Celaya 2014, Tlaltizapan, 2015 and Tlaltizapan, 2017)

**Table 2.** Descriptive statistics for kernel Zn concentration in (A) the ZAM panel and (B) DH populations grown in each environment, variance components and broad-and narrow sense heritabilities.

| Population | Population size | Location | Mean ± se ($\mu\text{g/g}$) | $\sigma_G^2$<sup>a</sup> | $\sigma_{GE}^2$<sup>a</sup> | $H^2$ |
| --- | --- | --- | --- | --- | --- | --- |
| ZAM panel | 923 | Agua Fria 2012 | 26.15 ± 0.15 | 12.04 | 2.42 | 0.85 |
| | | Celaya 2012 | 25.06 ± 0.14 | | | |
| | | Agua Fria 2013 | 29.53 ± 0.16 | | | |
|  |  | Across | **26.94 ± 0.10** |  |  |  |

**B**
| Population | Population size | Location | Mean ± se ($\mu\text{g/g}$) | $\hat{h}^2$ |
| --- | --- | --- | --- | --- |
| DH1 | 112 | Celaya 2014 | 25.38 ± 0.48 | 0.83 |
| | | Tlaltizapan 2015 | 24.01 ± 0.38 | |
| | | Tlaltizapan 2017 | 24.53 ± 0.37 | |
|  |  | Across | **24.65 ± 0.26** |  |
| DH2 | 143 | Celaya 2014 | 27.96 ± 0.39 | 0.76 |
| | | Tlaltizapan 2015 | 24.08 ± 0.33 | |
| | | Tlaltizapan 2017 | 24.64 ± 0.37 | |
|  |  | Across | **25.59 ± 0.22** |  |

Broad-sense heritability $H^2$ of Zn in each environment and across environments
Narrow-sense heritability $\hat{h}^2$ of Zn across environments
<sup>a</sup>variance due to genotypes $\sigma_G^2$ and the interaction between genotypes and the environment $\sigma_{GE}^2$ significant at $P < 0.001$

**Table 3.** Phenotypic correlation between environments for kernel Zn

|  | DH1 | DH 2 | ZAM Panel |
| --- | --- | --- | --- |
| <sup>a</sup>Env1 vs Env2 | 0.62 | 0.46 | 0.63 |
| <sup>a</sup>Env1 vs Env3 | 0.58 | 0.29 | 0.66 |
| <sup>a</sup>Env2 vs Env3 | 0.62 | 0.45 | 0.61 |

Phenotypic correlation coefficients were significant at $\alpha = 0.001$
<sup>a</sup>DH populations; Env 1, Env2 and Env 3=Celaya,2014, Tlaltizapan, 2017 and Tlaltizapan 2017, respectively.
<sup>a</sup>ZAM panel; Env 1, Env2 and Env 3= Agua Fria, 2012, Celaya, 2012 and Agua Fria 2013, respectively.

Principal component analysis for the ZAM panel suggested presence of a relatively diverse set of lines, and 452 principal components (PCs) were needed to explain 80% of the genotypes’ variance (Figures 3A and 3B). The first two principal components explained 3.85%of the total variance. For the DH populations first two eigenvectors separated them two groups (DH1 and DH2) and 56 principal components were needed to explain 80% of the genotypes’ variance (Figures 3C and 3D). The first two principal components explained 27.50% of the total variation for the DH populations.

**Figure 3.**
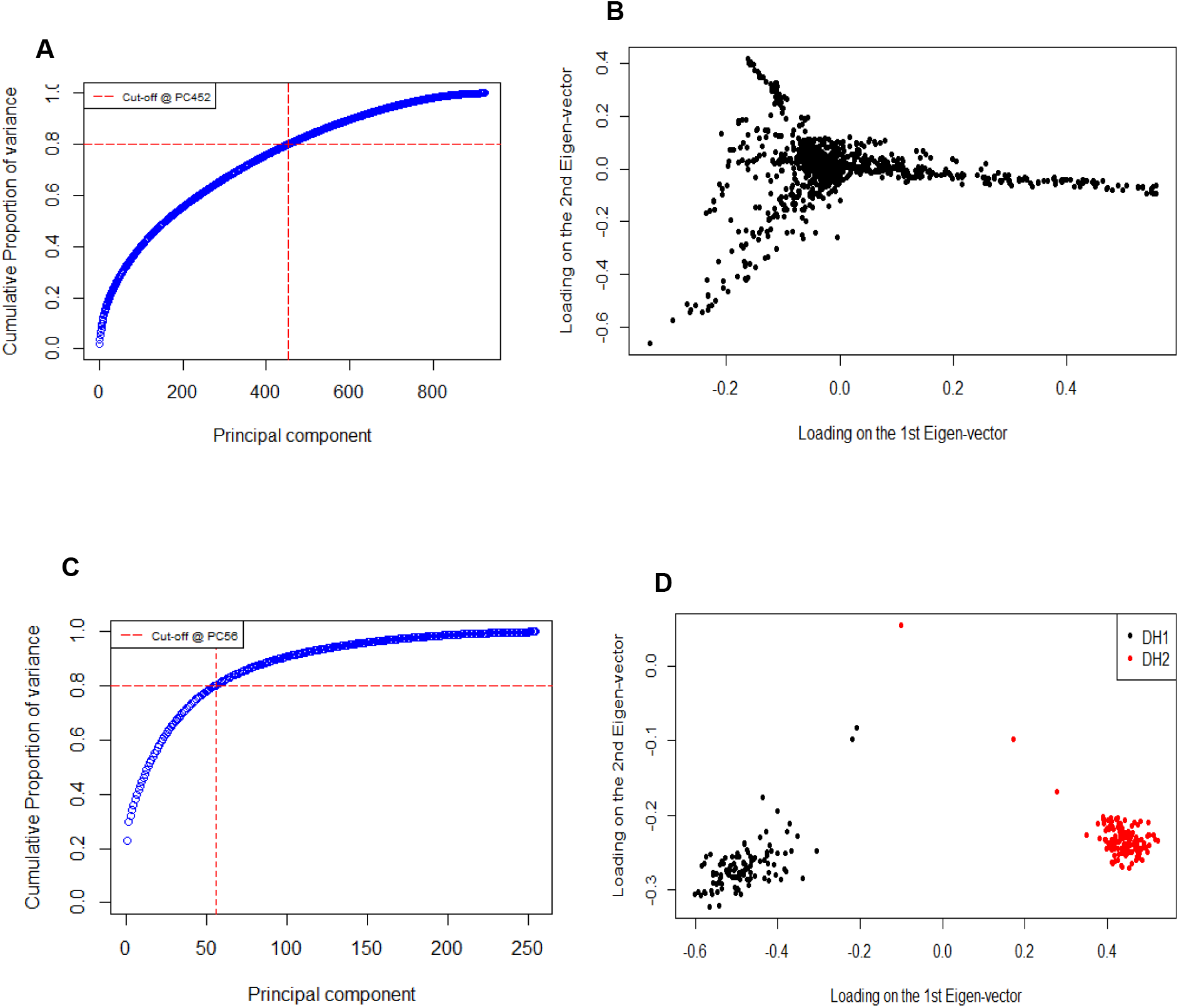
Scree plots (*A and C*) and loadings of the first two eigenvectors (*B and D*) of the covariance matrices derived from markers for the ZAM panel (A and B) and for the DH populations (*C and D*)

### Prediction ability in different populations

Cross-validated *r_MP_* values for kernel Zn were estimated for the ZAM panel and DH populations (Tables 4, 5 and 6). The average *r_MP_* values in CV1 were consistently lower than those in CV2, suggesting the importance of using information from correlated environments when predicting performance of inbred lines. The mean *r_MP_* values in CV1 and CV2 for the ZAM panel were 0.39 and 0.71, respectively (Table 4). For the DH populations, average *r_MP_* values were 0.53 for DH1-CV1, 0.44 for DH2-CV1 (Table 5), 0.70 for DH1-CV2 and 0.51 for DH2-CV2 (Table 6).

**Table 4.** Correlations (mean ± SD) between observed and genomic estimated breeding values for kernel Zn in the three environments for three GBLUP models for cross-validations CV1 and CV2 of the ZAM panel

| Population | Environment | Prediction accuracy in CV1 |  |  |
| --- | --- | --- | --- | --- |
|  |  | M1<sup>a</sup> | M2 | M3 |
| ZAM panel (923) | Agua Fria, 2012 | -0.01± 0.04 | 0.33 ± 0.01 | 0.34 ± 0.02 |
| | Celaya, 2012 | 0.004 ± 0.04 | 0.43 ± 0.01 | 0.47 ± 0.01 |
| | Agua Fria, 2013 | -0.001 ± 0.03 | 0.34 ± 0.01 | 0.35 ± 0.01 |
|  | **Average** | **-0.001± 0.03** | **0.37 ± 0.01** | **0.39 ± 0.01** |
| Population | Environment | Prediction accuracy in CV2 |  |  |
|  |  | <sup>a</sup>M1 | M2 | M3 |
| ZAM panel (923) | Agua Fria, 2012 | 0.71 ± 0.00 | 0.71 ± 0.00 | 0.72 ± 0.00 |
| | Celaya, 2012 | 0.64 ± 0.00 | 0.68 ± 0.00 | 0.72 ± 0.00 |
| | Agua Fria, 2013 | 0.67 ± 0.00 | 0.67 ± 0.00 | 0.69 ± 0.01 |
|  | **Average** | **0.67 ± 0.00** | **0.69 ± 0.00** | **0.71 ± 0.00** |

<sup>a</sup>Models: M1= Environment +Line; M2 = Environment + Line + Genomic; M3 = Environment + Line + Genomic + Genomic × Environment

**Table 5.** Correlations (mean ± SD) between observed and genomic estimated breeding values for Zn in the three environments for three GBLUP models for cross-validation CV1 of DH populations

| Population | Environment | Prediction accuracy in CV1 |  |  |
| --- | --- | --- | --- | --- |
|  |  | M1<sup>a</sup> | M2 | M3 |
| DH1 | Celaya, 2014 | -0.05 ± 0.10 | 0.52 ± 0.04 | 0.51 ± 0.04 |
| | Tlaltizapan, 2015 | -0.02 ± 0.12 | 0.52 ± 0.05 | 0.51 ± 0.05 |
| | Tlaltizapan, 2017 | -0.01 ± 0.10 | 0.56 ± 0.05 | 0.55 ± 0.05 |
|  | **Average** | **-0.03 ± 0.10** | **0.53 ± 0.04** | **0.52 ± 0.04** |
| DH2 | Celaya, 2014 | 0.05 ± 0.08 | 0.47 ± 0.03 | 0.50 ± 0.04 |
| | Tlaltizapan, 2015 | 0.03 ± 0.08 | 0.45 ± 0.03 | 0.45 ± 0.03 |
| | Tlaltizapan, 2017 | 0.04 ± 0.08 | 0.35 ± 0.03 | 0.35 ± 0.04 |
|  | **Average** | **0.04 ± 0.06** | **0.43 ± 0.03** | **0.44 ± 0.02** |

<sup>a</sup>Models: M1= Environment +Line; M2 = Environment + Line + Genomic; M3 = Environment + Line + Genomic + Genomic × Environment

**Table 6.** Correlations (mean ± SD) between observed and genomic estimated breeding values for Zn in the three environments for three GBLUP models for cross-validation CV2 of DH populations

| Population | Environment | Prediction accuracy in CV2 |  |  |
| --- | --- | --- | --- | --- |
|  |  | M1<sup>a</sup> | M2 | M3 |
| DH1 | Celaya, 2014 | 0.67 ± 0.02 | 0.68 ± 0.02 | 0.68 ± 0.03 |
| | Tlaltizapan, 2015 | 0.70 ± 0.02 | 0.71 ± 0.02 | 0.70 ± 0.02 |
| | Tlaltizapan, 2017 | 0.67 ± 0.02 | 0.70 ± 0.02 | 0.69 ± 0.02 |
|  | **Average** | **0.68 ± 0.01** | **0.70 ± 0.01** | **0.69 ± 0.01** |
| DH2 | Celaya, 2014 | 0.46 ± 0.016 | 0.53 ± 0.02 | 0.56 ± 0.02 |
| | Tlaltizapan, 2015 | 0.50 ± 0.020 | 0.55 ± 0.02 | 0.55 ± 0.02 |
| | Tlaltizapan, 2017 | 0.40 ± 0.023 | 0.43 ± 0.02 | 0.43 ± 0.02 |
|  | **Average** | **0.45 ± 0.02** | **0.50 ± 0.01** | **0.51 ± 0.01** |

<sup>a</sup>Models: M1= Environment +Line; M2 = Environment + Line + Genomic; M3 = Environment + Line + Genomic + Genomic × Environment

In the ZAM panel, the highest values in CV1 (0.47) and CV2 (0.72) were obtained in Celaya and Agua Fria 2012 (Table 4). For the bi-parental populations, both under CV1 and CV2, higher *r_MP_* values were observed for DH1 compared to DH2. The highest values in CV1 (0.56) and CV2 (0.71) were observed in Tlaltizapan 2017 and 2015, all for DH1 (Tables 5 and 6). The consistently higher *r_MP_* values in CV1 and CV2 of DH1 could be attributed to the higher (0.58 to 0.62) correlation values between environments (Table 3).

### Prediction ability of different models

Comparing the *r_MP_* values obtained from each model, M1 had the lowest (−0.001, −0.03 and 0.04) accuracies in CV1 for the ZAM panel and DH populations (Tables 4 and 5). Those values were improved in CV2 because the predictions benefited from previous records (collected from other environments) of lines whose Zn values were being predicted. When M1 was expanded to M2 by adding the main effects of markers, the *r_MP_* values at each environment and across environments were increased. For example, in CV1, M2, >100-fold increase in *r_MP_* values were observed for the ZAM panel and DH populations, and in CV2, M2, average *r_MP_* values increased by 2.98%, 2.94% and 11.11% for the ZAM panel, DH1 and DH2, respectively (Tables 4, 5 and 6).

The multi-environment model (M3), which includes the interaction between markers (genomic) and the environment (*Eg_ij_*) gave higher prediction accuracy than single-environment models (M1 and M2). In CV1, mean *r_MP_* values increased from 0.37 (M2) to 0.39 (M3) for the ZAM panel and from 0.43 (M2) to 0.44 for DH2 (Tables 4 and 5). Similar trends were observed in CV2 for the ZAM panel and DH2 (Tables 4 and 6). However, in both CV1 and CV2 of DH1, incorporating *Eg_ij_* did not improve *r_MP_* values for Zn (Tables 5 and 6). For CV1, M3, *r_MP_* values for Zn in individual environments ranged from 0.34 to 0.47 for the ZAM panel (Table 4), 0.51 to 0.55 for DH1 and 0.35 to 0.50 for DH2 (Table 5). For CV2, M3, those values ranged from 0.69 to 0.72 for the ZAM panel, 0.68 to 0.70 for DH1 and 0.43 to 0.56 for DH2 (Tables 4, 5 and 6).

## DISCUSSION

Overall, moderate to high prediction ability values for kernel Zn were observed for the ZAM panel and DH populations. This could be attributed to the heritabilities observed for kernel Zn (Tables 2A and 2B). Similar observations were reported for Zn concentration in wheat (Velu *et al*. 2016; Manickavelu *et al*. 2017). High quality predictions with high accuracy for GS programs are expected for traits with moderate to higher heritability estimates (Combs and Bernardo 2013; Lian *et al*. 2014; Muranty *et al*. 2015; Saint Pierre *et al*. 2016; Manickavelu *et al*. 2017; Zhang *et al*. 2017b, 2019; Arojju *et al*. 2019). Consistent with a study on Zn and iron (Fe) concentration in spring wheat, the prediction accuracies in this study are sufficient to discard at least 50% of the inbreds with low-Zn concentration (Velu *et al*. 2016).

Data from both bi-parental populations and diverse collection of inbreds have been used for GS and cross-validation (CV) experiments have shown that prediction accuracies could also be affected by the relatedness between training and prediction sets (Habier *et al*. 2007; de Roos *et al*. 2009; Asoro *et al*. 2011; Daetwyler *et al*. 2013; Cericola *et al*. 2017; Crossa *et al*. 2017). In this study, average predicted accuracies were higher for CV1 of the bi-parental populations (0.53 for DH1 and 0.44 for DH2) compared to the ZAM panel (0.39). Higher predicted values in CV1 of the DH populations could be attributed to the closer relationship between DH lines in the training and prediction sets, maximum linkage disequilibrium (LD) between a marker and a QTL, and controlled population structure (Bernardo and Yu 2007; Albrecht *et al*. 2011; Zhang *et al*. 2015). In collections of diverse inbreds, prediction accuracy may depend on the ancestral relationships between the lines. So, in experiments using such collections of lines, prediction accuracies have been more variable than accuracies achieved using bi-parental populations (Spindel and McCouch 2016).

Cross-validation (CV) schemes are used in genomic prediction to estimate the accuracy with which predictions for different traits and environments can be made (Burgueño *et al*. 2012; Zhang *et al*. 2015; Saint Pierre *et al*. 2016; Velu *et al*. 2016; Sukumaran *et al*. 2017a, 2017b; Monteverde *et al*. 2018; Roorkiwal *et al*. 2018). In this study, two CV schemes (CV1-predicting the performance of newly developed lines, and CV2-predicting the performance of lines that have been evaluated in some environments, but not in others) were used. The utility of these schemes indicated that prediction values for newly developed lines (CV1) were generally lower (0.39 for the ZAM panel, 0.53 for DH1 and 0.44 for DH2) than the values for lines which have been evaluated in different but correlated environments (CV2; 0.71, 0.70 and 0.51 for the ZAM panel, DH1 and DH2, respectively). Such observations indicate the importance of using information from correlated environments when predicting the performance of inbred lines. However, selection of new lines without field testing, as simulated in CV1 allows shortening of the generation interval (cycle time) by replacing the time-intensive phenotypic evaluation for Zn with genomic-estimated breeding values. But, the quality of prediction accuracy may be lower such that the annual rate of genetic progress in a GS program is compromised (Burgueño *et al*.2012). So, the ultimate decision of how a breeding scheme should be structured could depend on the compromise between the desired prediction accuracy and the generation interval (Burgueño *et al*. 2012).

Genotype by environment interaction is an important factor affecting kernel Zn concentration in maize and genomic prediction models that incorporate G x E may enhance the potential of GS for biofortification breeding. For different crop species and traits, genomic prediction models which incorporated G x E achieved higher prediction accuracies in both CV1 and CV2 schemes relative to models which did not include G x E (Burgueño *et al*. 2012; Guo *et al*. 2013; Jarquín *et al*. 2014; Lopez-Cruz *et al*. 2015; Zhang *et al*. 2015; Monteverde *et al*. 2018). In this study, the impact of modeling G x E variance structures for multi-environment trials was investigated and results indicated that the average predicted values from M3 (G x E model) were higher (0.39 and 0.44 for CV1 and 0.71 and 0.51 for CV2) than the values from M2 (non-G x E; 0.37 and 0.43 for CV1-M2, 0.69 and 0.50 for CV2-M2) for the ZAM panel and DH2. These findings agree with those reported on Zn concentration in wheat (Velu *et al*. 2016), providing evidence that incorporating G x E in GS models can enhance their power and suitability for improving maize for kernel Zn concentration. Conversely, the average predicted values for CV1 and CV2 of DH1 were higher in M2 (0.53 and 0.70) than in M3 (0.53 and 0.69). Except for differences in population size (112 lines vs 143 lines), this was unexpected since DH1 and DH2 were grown in the same environments.

The gains in prediction accuracies for the GS model that accounted for G x E were dependent on the correlation between environments and CV method used. In this study, the phenotypic correlations between environments were all positive (ranging from 0.58 to 0.62 for DH1, 0.29 to 0.46 for DH2 and 0.61 to 0.66 for the ZAM panel). Such correlations can be exploited using multi-environment models to derive predictions that use information from across both the lines and environments (Burgueño *et al*. 2012). For instance, although the phenotypic correlations between environments for DH2 were positive (0.29 to 0.46), the lowest average prediction value (0.51) for CV2 was observed for this population. This was expected because CV2 uses phenotypic information from genotypes which have already been tested; hence, effectively exploiting the correlations between environments (Burgueño *et al*. 2012; Jarquín *et al*. 2014; Crossa *et al*. 2015; Pérez-Rodríguez *et al*. 2015; Saint Pierre *et al*. 2016; Monteverde *et al*. 2018). However, for CV1, the information between environments could only be accounted for through the genomic relationship matrix (Monteverde *et al*. 2018). Hence, the gains in CV1 may likely attribute to more accurate estimate of environment-specific marker effects (Guo *et al*. 2013). In contrast, when multiple environments are weakly correlated, prediction accuracies from across environment analyses can be negatively affected relative to prediction accuracies within environments (Bentley *et al*. 2014; Wang *et al*. 2014; Spindel and McCouch 2016). Thus, before designing a GS experiment, identifying correlated environments where environments can differ in terms of site, year or season in which data were collected is of great interest (Spindel and McCouch 2016).

The ability to predict kernel Zn concentration using high-throughput SNP markers including G x E interactions creates an opportunity for efficiently enhancing Zn concentration in maize breeding programs. For instance, during early generations of a breeding program, GS can be utilized to identify genotypes with favorable alleles when numbers of progenies and families are large. This could potentially reduce the resource-intensive evaluation process and advancement of false-positive progenies (Velu *et al*. 2016). Coupled with advances in technologies for assessing Zn, plant scientists can more rapidly measure Zn concentration in maize kernels using the energy dispersive x-ray fluorescence (XRF) assays (Guild *et al*. 2017). Thus, with more validations and model refinements, GS can potentially accelerate the breeding process to enhance Zn concentration in maize for a wider range of environments.

## CONCLUSION

The moderate to high prediction accuracies reported in this study shows that GS can be used in maize breeding to improve kernel Zn concentration. Assuming two possible seasons of Zn evaluation per year, the predicted genetic gains can be estimated from prediction accuracies and genetic variances of the training populations. The genetic variances for the ZAM panel, DH1 and DH2 were 12.38, 12.20 and 14.88, and prediction accuracies were 0.71, 0.70 and 0.51, respectively. If the inbreds in each predicted population are ranked based on their predicted Zn values and the top 10% selected, then their expected average Zn values can be estimated from the proportion of inbreds selected, their respective training population genetic variances, prediction accuracies and the time interval for evaluating the lines. With reference to this, the expected average values of Zn are approximately 31 μg/g for the ZAM panel, 30 μg/g for DH1 and 27 μg/g for DH2. These averages are higher than the averages of the respective training populations (~27 μg/g for the ZAM panel, ~25 μg/g for DH1 and ~26 μg/g for DH2) suggesting that the prediction accuracies achieved are sufficient to select at least 10% of the predicted inbreds with higher Zn concentration.

The prediction accuracies were of lower quality when genomic predictions were conducted across populations. When the ZAM panel was used as the training population, prediction accuracies for DH1, DH2 and DH1+DH2 were 0.15, −0.10 and 0.09, respectively. When DH1 and DH2 were used as a training and prediction set for each other, prediction accuracies were 0.08 and 0.16 (Unpublished data). These prediction accuracies are considerably lower than those reported in this study and the differences may be attributed to: (i) weak genetic relationships between the training and prediction population sets and (ii) different methods of analysis because the prediction accuracies reported in this study were partly achieved by modelling the random-effects environment structure to account for G x E while for the unpublished data, the random-effects environment structure of G x E was not included.

This study also showed that higher prediction accuracies can be achieved when some of the lines are predicted using previous information about them collected from correlated environments. The multi-environment model (M3) which included the interaction between markers, and the environment gave higher prediction accuracy both in CV1 and CV2 for the association panel and DH2 compared with the models which only included main effects (M1 and M2) indicating the importance of accounting for G x E in genomic prediction.

## ACKNOWLEDGEMENTS

We thank HarvestPlus through CIMMYT and the R.F. Baker Center for Plant Breeding, Department of Agronomy at Iowa State University for making this research possible. We also would like to thank, Andrea Cruz-Morales, Mayolo Leyva, Balfre Tellez-Noriega and Adolfo Basilio for managing the trials and collecting field data. We are grateful to Aldo Rosales and staff of the Maize Grain Quality laboratory at CIMMYT for carrying out the micronutrient analyses.

## LITERATURE CITED

Agrawal, P. K., S. K. Jaiswal, B. M. Prasanna, F. Hossain, S. Saha et al., 2012 Genetic variability and stability for kernel iron and zinc concentration in maize (Zea mays L.) genotypes. Indian J. Genet. Plant Breed. 72: 421–428.

Ahmadi, M., W. J. Wiebold, J. E. Beuerlein, D. J. Eckert, and J. Schoper, 1993 Agronomic Practices that Affect Corn Kernel Characteristics. Agron. J. 85: 615–619.

Albrecht, T., V. Wimmer, H. J. Auinger, M. Erbe, C. Knaak et al., 2011 Genome-based prediction of testcross values in maize. Theor. Appl. Genet. 123: 339–350.

Alvarado, G., M. López, M. Vargas, A. Pacheco, F. Rodríguez et al., 2016 META-R (Multi Environment Trial Analysis with R for Windows) Version 6.04.

Arojju, S. K., M. Cao, M. Z. Zulfi Jahufer, B. A. Barrett, and M. J. Faville, 2019 Genomic Predictive Ability for Foliar Nutritive Traits in Perennial Ryegrass. G3 10: 695–708.

Asoro, F. G., M. A. Newell, W. D. Beavis, M. P. Scott, and J.-L. Jannink, 2011 Accuracy and Training Population Design for Genomic Selection on Quantitative Traits in Elite North American Oats. Plant Genome 4: 132–144.

Bänziger, M., and J. Long, 2000 The potential for increasing the iron and zinc density of maize through plant-breeding. Food Nutr. Bull. 21: 397–400.

Baxter, I. R., J. L. Gustin, A. M. Settles, and O. A. Hoekenga, 2013 Ionomic characterization of maize kernels in the intermated B73 x Mo17 population. Crop Sci. 53: 208–220.

Bentley, A. R., M. Scutari, N. Gosman, S. Faure, F. Bedford et al., 2014 Applying association mapping and genomic selection to the dissection of key traits in elite European wheat. Theor. Appl. Genet. 127: 2619–2633.

Bernardo, R., and J. Yu, 2007 Prospects for genomewide selection for quantitative traits in maize. Crop Sci. 47: 1082–1090.

Bouis, H. E., and A. Saltzman, 2017 Improving nutrition through biofortification: A review of evidence from HarvestPlus, 2003 through 2016. Glob. Food Sec. 12: 49–58.

Brkic, I., D. Simic, Z. Zdunic, A. Jambrovic, T. Ledencan et al., 2004 Genotypic variability of micronutrient element concentrations in maize kernels. Cereal Res. Commun. 32: 107–112.

Burgueño, J., G. de los Campos, K. Weigel, and J. Crossa, 2012 Genomic prediction of breeding values when modeling genotype × environment interaction using pedigree and dense molecular markers. Crop Sci. 52: 707–719.

Cericola, F., A. Jahoor, J. Orabi, J. R. Andersen, L. L. Janss et al., 2017 Optimizing Training Population Size and Genotyping Strategy for Genomic Prediction Using Association Study Results and Pedigree Information. A Case of Study in Advanced Wheat Breeding Lines. PLoS One 12: e0169606.

Chakraborti, M., B. M. Prasanna, F. Hossain, S. Mazumdar, A. M. Singh et al., 2011 Identification of kernel iron- and zinc-rich maize inbreds and analysis of genetic diversity using microsatellite markers. J. Plant Biochem. Biotechnol. 20: 224–233.

Chakraborti, M., B. M. Prasanna, F. Hossain, A. M. Singh, and S. K. Guleria, 2009 Genetic evaluation of kernel Fe and Zn concentrations and yield performance of selected Maize (Zea mays L.) genotypes. Range Manag. Agrofor. 30: 109–114.

CIMMYT, 2005 Laboratory Protocols: CIMMYT Applied Molecular Genetics Laboratory.

Combs, E., and R. Bernardo, 2013 Accuracy of genomewide selection for different traits with constant population size, heritability, and number of markers. Plant Genome 6: 1–7.

core Team, R., 2018 R: A Language and Environment for Statistical Computing. R Found. Stat. Comput. Vienna, Austria.

Crossa, J., Y. Beyene, S. Kassa, P. Pérez, J. M. Hickey et al., 2013 Genomic prediction in maize breeding populations with genotyping-by-sequencing. G3 3: 1903–1926.

Crossa, J., G. de los Campos, M. Maccaferri, R. Tuberosa, J. Burgueño et al., 2015 Extending the marker × Environment interaction model for genomic-enabled prediction and genome-wide association analysis in durum wheat. Crop Sci. 56: 2193–2209.

Crossa, J., G. de los Campos, P. Pérez, D. Gianola, J. Burgueño et al., 2010 Prediction of genetic values of quantitative traits in plant breeding using pedigree and molecular markers. Genetics 186: 713–724.

Crossa, J., P. Pérez-Rodríguez, J. Cuevas, O. Montesinos-López, D. Jarquín et al., 2017 Genomic Selection in Plant Breeding: Methods, Models, and Perspectives. Trends Plant Sci. 22:961–975.

Crossa, J., P. Pérez, J. Hickey, J. Burgueño, L. Ornella et al., 2014 Genomic prediction in CIMMYT maize and wheat breeding programs. Heredity (Edinb). 112: 48–60.

Cuevas, J., J. Crossa, O. A. Montesinos-López, J. Burgueño, P. Pérez-Rodríguez et al., 2017 Bayesian genomic prediction with genotype × environment interaction kernel models. G3 7: 41–53.

Cuevas, J., J. Crossa, V. Soberanis, S. Pérez-Elizalde, P. Pérez-Rodríguez et al., 2016 Genomic prediction of genotype × environment interaction kernel regression models. Plant Genome 9: 1–20.

Daetwyler, H. D., M. P. L. Calus, R. Pong-Wong, G. de los Campos, and J. M. Hickey, 2013 Genomic prediction in animals and plants: Simulation of data, validation, reporting, and benchmarking. Genetics 193: 347–365.

Elshire, R. J., J. C. Glaubitz, Q. Sun, J. A. Poland, K. Kawamoto et al., 2011 A robust, simple genotyping-by-sequencing (GBS) approach for high diversity species. PLoS One 6: e19379.

Gannon, B. M., K. V Pixley, and S. A. Tanumihardjo, 2017 Maize Milling Method Affects Growth and Zinc Status but Not Provitamin A Carotenoid Bioefficacy in Male Mongolian Gerbils. J. Nutr. 147: 337–345.

Glaubitz, J. C., T. M. Casstevens, F. Lu, J. Harriman, R. J. Elshire et al., 2014 TASSEL-GBS: A high capacity genotyping by sequencing analysis pipeline. PLoS One 9: e90346.

Govindan, V., 2011 Breeding for enhanced Zinc and Iron concentration in CIMMYT spring wheat germplasm. Czech J. Genet. Plant Breed. 47: 174–177.

Guild, G. E., N. G. Paltridge, M. S. Andersson, and J. C. R. Stangoulis, 2017 An energy-dispersive X-ray fluorescence method for analysing Fe and Zn in common bean, maize and cowpea biofortification programs. Plant Soil 419: 457–466.

Guleria, S. K., R. K. Chahota, P. Kumar, A. Kumar, B. M. Prasanna et al., 2013 Analysis of genetic variability and genotype x year interactions on kernel zinc concentration in selected Indian and exotic maize (Zea mays) genotypes. Indian J. Agric. Sci. 83: 836–841.

Guo, Z., D. M. Tucker, D. Wang, C. J. Basten, E. Ersoz et al., 2013 Accuracy of across-environment genome-wide prediction in maize nested association mapping populations. G3 3: 263–272.

Habier, D., R. L. Fernando, and J. C. M. Dekkers, 2007 The impact of genetic relationship information on genome-assisted breeding values. Genetics 177: 2383–2397.

Heslot, N., J.-L. Jannink, and M. E. Sorrells, 2015 Perspectives for Genomic Selection Applications and Research in Plants. Crop Sci. 55: 1–12.

Hindu, V., N. Palacios-Rojas, R. Babu, W. B. Suwarno, Z. Rashid et al., 2018 Identification and validation of genomic regions influencing kernel zinc and iron in maize. Theor. Appl. Genet. 131: 1443–1457.

Jarquín, D., J. Crossa, X. Lacaze, P. Du Cheyron, J. Daucourt et al., 2014 A reaction norm model for genomic selection using high-dimensional genomic and environmental data. Theor. Appl. Genet. 127: 595–607.

Jarquín, D., C. Lemes da Silva, R. C. Gaynor, J. Poland, A. Fritz et al., 2017 Increasing Genomic-Enabled Prediction Accuracy by Modeling Genotype × Environment Interactions in Kansas Wheat. Plant Genome 10: 1–15.

Jin, T., J. Zhou, J. Chen, L. Zhu, Y. Zhao et al., 2013 The genetic architecture of zinc and iron content in maize grains as revealed by QTL mapping and meta-analysis. Breed. Sci. 63: 317–324.

Lian, L., A. Jacobson, S. Zhong, and R. Bernardo, 2014 Genomewide prediction accuracy within 969 maize biparental populations. Crop Sci. 54: 1514–1522.

Listman, M., 2019 Biofortified maize and wheat can improve diets and health, new study shows. Int. Maize Wheat Improv. Cent.

Liu, X., H. Wang, H. Wang, Z. Guo, X. Xu et al., 2018 Factors affecting genomic selection revealed by empirical evidence in maize. Crop J. 6: 341–352.

Long, J. K., M. Bänziger, and M. E. Smith, 2004 Diallel analysis of grain iron and zinc density in southern African-adapted maize inbreds. Crop Sci. 44: 2019–2026.

Lopez-Cruz, M., J. Crossa, D. Bonnett, S. Dreisigacker, J. Poland et al., 2015 Increased prediction accuracy in wheat breeding trials using a marker × environment interaction genomic selection model. G3 5: 569–582.

de los Campos, G., H. Naya, D. Gianola, J. Crossa, A. Legarra et al., 2009 Predicting quantitative traits with regression models for dense molecular markers and pedigree. Genetics 182: 375–385.

de los Campos, G., P. Pérez, A. I. Vazquez, and J. Crossa, 2013 Genome-enabled prediction using the BLR (Bayesian Linear Regression) R-package, pp. 229–320 in Genome-Wide Association Studies and Genomic Prediction. Methods in Molecular Biology (Methods and Protocols), edited by C. Gondro, J. van der Werf, and B. Hayes. Humana Press, Totowa, NJ.

Manickavelu, A., T. Hattori, S. Yamaoka, K. Yoshimura, Y. Kondou et al., 2017 Genetic nature of elemental contents in wheat grains and its genomic prediction: Toward the effective use of wheat landraces from Afghanistan. PLoS One 12: e0169416.

Menkir, A., 2008 Genetic variation for grain mineral content in tropical-adapted maize inbred lines. Food Chem. 110: 454–464.

Meuwissen, T. H. E., B. J. Hayes, and M. E. Goddard, 2001 Prediction of total genetic value using genome-wide dense marker maps. Genetics 157: 1819–1829.

Misra, B. K., R. K. Sharma, and S. Nagarajan, 2004 Plant breeding: A component of public health strategy. Curr. Sci. Assoc. 86: 1210–1215.

Monteverde, E., J. E. Rosas, P. Blanco, F. Pérez de Vida, V. Bonnecarrère et al., 2018 Multienvironment models increase prediction accuracy of complex traits in advanced breeding lines of rice. Crop Sci. 58: 1519–1530.

Muranty, H., M. Troggio, I. Ben Sadok, M. Al Rifai, A. Auwerkerken et al., 2015 Accuracy and responses of genomic selection on key traits in apple breeding. Hortic. Res. 2: 1–12.

Oikeh, S. O., A. Menkir, Bussie Maziya-Dixon, Ross Welch, and R. P. Glahn, 2003 Assessment of Concentrations of Iron and Zinc and Bioavailable Iron in Grains of Early-Maturing Tropical Maize Varieties. J. Agric. Food Chem. 3688–3694.

Oikeh, S. O., a. Menkir, B. Maziya-Dixon, R. M. Welch, R. P. Glahn et al., 2004 Environmental stability of iron and zinc concentrations in grain of elite early-maturing tropical maize genotypes grown under field conditions. J. Agric. Sci. 142: 543–551.

Palacios-Rojas, N., 2018 Calidad nutricional e industrial de Maíz: Laboratorio de Calidad Nutricional de Maíz. Mexico.

Pérez-Rodríguez, P., J. Crossa, K. Bondalapati, G. De Meyer, F. Pita et al., 2015 A pedigree-based reaction norm model for prediction of cotton yield in multienvironment trials. Crop Sci. 55: 1143–1151.

Pérez-Rodríguez, P., D. Gianola, J. M. González-Camacho, J. Crossa, Y. Manès et al., 2012 Comparison between linear and non-parametric regression models for genome-enabled prediction in wheat. G3 2: 1595–1605.

Pérez, P., and G. de los Campos, 2014 Genome-wide regression and prediction with the BGLR statistical package. Genetics 198: 483–495.

Saint Pierre, C., J. Burgueño, J. Crossa, G. Fuentes Dávila, P. Figueroa López et al., 2016 Genomic prediction models for grain yield of spring bread wheat in diverse agro-ecological zones. Sci. Rep. 6: 1–11.

Prasanna, B. M., S. Mazumdar, M. Chakraborti, F. Hossain, and K. M. Manjaiah, 2011 Genetic variability and genotype × environment interactions for kernel iron and zinc concentrations in maize (Zea mays) genotypes. Indian J. Agric. Sci. 81: 704–711.

Pszczola, M., T. Strabel, H. A. Mulder, and M. P. L. Calus, 2012 Reliability of direct genomic values for animals with different relationships within and to the reference population. J. Dairy Sci. 95: 389–400.

Qin, H., Y. Cai, Z. Liu, G. Wang, J. Wang et al., 2012 Identification of QTL for zinc and iron concentration in maize kernel and cob. Euphytica 187: 345–358.

Roorkiwal, M., D. Jarquin, M. K. Singh, P. M. Gaur, C. Bharadwaj et al., 2018 Genomic-enabled prediction models using multi-environment trials to estimate the effect of genotype × environment interaction on prediction accuracy in chickpea. Sci. Rep. 8: 1–11.

de Roos, A. P. W., B. J. Hayes, and M. E. Goddard, 2009 Reliability of genomic predictions across multiple populations. Genetics 183: 1545–1553.

Ŝimić, D., S. Mladenović Drinić, Z. Zdunić, A. Jambrović, T. Ledenan et al., 2012 Quantitative trait loci for biofortification traits in maize grain. J. Hered. 103: 47–54.

Spindel, J. E., and S. R. McCouch, 2016 When more is better: how data sharing would accelerate genomic selection of crop plants. New Phytol. 212: 814–826.

Stein, A. J., 2010 Global impacts of human mineral malnutrition. Plant Soil 335: 133–154.

Sukumaran, S., J. Crossa, D. Jarquin, M. Lopes, and M. P. Reynolds, 2017a Genomic prediction with pedigree and genotype × environment interaction in spring wheat grown in South and West Asia, North Africa, and Mexico. G3 7: 481–495.

Sukumaran, S., J. Crossa, D. Jarquín, and M. Reynolds, 2017b Pedigree-based prediction models with genotype × environment interaction in multienvironment trials of CIMMYT wheat. Crop Sci. 57: 1865–1880.

VanRaden, P. M., 2008 Efficient methods to compute genomic predictions. J. Dairy Sci. 91: 4414–4423.

Velu, G., J. Crossa, R. P. Singh, Y. Hao, S. Dreisigacker et al., 2016 Genomic prediction for grain zinc and iron concentrations in spring wheat. Theor. Appl. Genet. 129: 1595–1605.

Wang, Y., M. F. Mette, T. Miedaner, M. Gottwald, P. Wilde et al., 2014 The accuracy of prediction of genomic selection in elite hybrid rye populations surpasses the accuracy of marker-assisted selection and is equally augmented by multiple field evaluation locations and test years. BMC Genomics 15: 1–15.

Welch, R. M., and R. D. Graham, 2004 Breeding for micronutrients in staple food crops from a human nutrition perspective. J. Exp. Bot. 55: 353–364.

Zhang, H., J. Liu, T. Jin, Y. Huang, J. Chen et al., 2017a Identification of quantitative trait locus and prediction of candidate genes for grain mineral concentration in maize across multiple environments. Euphytica 213: 1–16.

Zhang, X., P. Pérez-Rodríguez, K. Semagn, Y. Beyene, R. Babu et al., 2015 Genomic prediction in biparental tropical maize populations in water-stressed and well-watered environments using low-density and GBS SNPs. Heredity (Edinb). 114: 291–299.

Zhang, A., H. Wang, Y. Beyene, K. Semagn, Y. Liu et al., 2017b Effect of Trait Heritability, Training Population Size and Marker Density on Genomic Prediction Accuracy Estimation in 22 bi-parental Tropical Maize Populations. Front. Plant Sci. 8: 1–12.

Zhang, H., L. Yin, M. Wang, X. Yuan, and X. Liu, 2019 Factors affecting the accuracy of genomic selection for agricultural economic traits in maize, cattle, and pig populations. Front. Genet. 10: 1–10.

Zhao, Y., M. Gowda, W. Liu, T. Würschum, H. P. Maurer et al., 2012 Accuracy of genomic selection in European maize elite breeding populations. Theor. Appl. Genet. 124: 769–776.

Zhou, J.-F., Y.-Q. Huang, Z.-Z. Liu, J.-T. Chen, L.-Y. Zhu et al., 2010 Genetic Analysis and QTL Mapping of Zinc, Iron, Copper and Manganese Contents in Maize Seed. J. Plant Genet. Resour. 11: 593–595.

